# Phenotypic Resistance of Ciprofloxacin and Azithromycin Resistant *Campylobacter* Isolates to an Extended Panel of Antibiotics

**DOI:** 10.64898/2026.02.06.704332

**Authors:** Lucero Romaina_Cachique, Francesca Schiaffino, Maribel Paredes_Olortegui, Katia Manzanares, Tackeshy Pinedo, Alejandra Castro, Juana Najarro, Kerry K. Cooper, Evangelos Mourkas, Ben Pascoe, Pablo Peñataro_Yori, Craig T. Parker, Margaret N. Kosek

**Author notes:** **Address correspondence to:** Margaret N. Kosek; University of Virginia, Division of Infectious Diseases, International Health, and Public Health Sciences; Address: 345 Crispell Dr, Rm 2525, Charlottesville, VA.

## Abstract

In this study we explored phenotypic resistance using traditional Kirby-Bauer methods in human and animal derived *Campylobacter* isolates that were concurrently resistant to both azithromycin and ciprofloxacin to an expanded panel of antimicrobials, including clindamycin, fosfomycin, ampicillin sulbactam and tigecycline. Out of 236 *Campylobacter* isolates, over 85% of *C. jejuni* and *C. coli* were resistant to clindamycin, over 60% were resistant to ampicillin sulbactam and over 30% to fosfomycin. Less than 2% of isolates were resistant to tigecycline and there was no observed resistance to Imipenem.

## Introduction

*Campylobacter* infections are a leading global cause of bacterial enteritis^1-3^. Additionally, *Campylobacter* are among the foremost pathogens associated with foodborne illnesses, traveler’s diarrhea and animal contact in high income countries, causing 1.9 million illnesses and up to 13,000 hospitalizations in the United States.^4-8^ Although antimicrobials are not advised for treatment for uncomplicated gastroenteritis, antimicrobial therapy is advised for patients with severe symptoms, dysentery, and persistent infections, as well as in patients who are immunocompromised or pregnant.^9^

Fluoroquinolones and azithromycin (macrolides) are the two most effective oral antimicrobial treatments for campylobacteriosis in sensitive strains, and have been shown to shorten the duration of illness in clinical trials.^10^ Macrolide resistance in *Campylobacter jejuni* has remained stable in the United States between 2013 and 2019, while quinolone resistance has shown an increasing trend, according to National Antimicrobial Resistance Monitoring System (NARMS)^11^. However, a rising occurrence of *C. jejuni* and *C. coli* isolates resistant macrolides has been evidenced in other scenarios such as the outbreak in the U.S. linked to puppies, making it clear that there is a need to evaluate additional antibiotics to be prepared for similar future outbreaks and progressive antimicrobial resistance (AMR) in campylobacteriosis.^12-14^ This study attempted to define possible alternative therapies among strains resistant to both fluoroquinolones and azithromycin.

The increase in AMR among *C. jejuni* and *C. coli* has occurred globally and is likely result of the extensive use of antimicrobials in livestock animals, especially in regions with poor or minimal enforcement of restrictions in the use of agroindustry. In Peru, prevalence trends of AMR in *C. jejuni* and *C. coli* have primarily focused on resistances to fluoroquinolones and macrolides in humans given their importance in treatment of human illnesses.^15-17^ Resistance to fluoroquinolones has increased dramatically in the Peru from 2001 to 2010 ^15,16^ and remains at higher levels through 2022.^17,18^ Genomic of analysis *C. jejuni* and *C. coli* isolates in Iquitos, Peru has identified specific chromosomal mutations for resistance, including a *gyrA* Thr86Ile mutation associated with fluoroquinolone resistance, the 23S rRNA mutations associated with macrolide resistance, and a modification of cmeABC efflux pump (RE-*cmeAB*C) that confers multi-drug resistance (defined as non-susceptibility to at least one antibiotic from three or more antimicrobial clases).^18-23^

Given the lack of clearly defined additional options to treat clinical cases of campylobacteriosis demonstrating resistance to both fluoroquinolones and azithromycin, there is a need to evaluate additional antibiotics that are commonly used to treat gastrointestinal infections, particularly oral therapies. We explored *in vitro* phenotypic resistance patterns in a broad collection of *C. jejuni* and *C. coli* isolates derived from human and animal fecal samples to an expanded set of antibiotics that have not been widely evaluated.

## Methods

Infant fecal samples, animal fecal samples and meat during a period of an ongoing cohort study conducted in the Peruvian Amazon from 2021-2025. Human fecal samples were collected monthly or every time the child had diarrhea. Animal fecal samples were derived from a nested case-control study associated with the same cohort, or from concurrent surveillance conducted throughout the duration of the cohort study were livestock (chickens, pigs, cows), household animals and chicken meat were assessed for the presence of *Campylobacter*. The study was approved by the Institutional Review Boards of Asociacion Benefica Prisma, the University of Virginia and Universidad Peruana Cayetano Heredia. Written informed consent to participate in the study was obtained from the parents or legal guardians of children. Participants consented for further use of biological specimens.

Human and animal fecal samples were cultured using Columbia Blood Agar Base (Oxoid, Thermo Fisher Scientific, Waltham, MAS) supplemented with 5% lysed horse blood and an S-pack filter of 0.45uM and 47mm in diameter (Merck Millipore, Burlington, MAS), in microaerophilic conditions (1% O_2_ + 10% CO_2_ + 10% H_2_ + balance N_2_) at 37°C. Colonies compatible with *Campylobacter* morphology were further confirmed as *Campylobacter* spp. Or *C. jejuni* / *C. coli* using a duplex qPCR targeting the 16S rRNA and a rpsS / rpsK gene in *Campylobacter*.^24,25^ Only isolates for which a species (*C. jejuni* or *C. coli*) could be assigned were included in the final data set for analysis.

Phenotypic antimicrobial susceptibility testing was performed using standard disc-diffusion testing.^17^ Resistance to the following antibiotics was tested: ciprofloxacin (CIP), erythromycin (ERY), azithromycin (AZM), tetracycline (TET), gentamicin (GEN), amoxicillin and clavulanic acid (AMC), chloramphenicol (CHL) and imipenem (IMP). An extended panel including clindamycin (CLI), fosfomycin (FOSF), ampicillin sulbactam (AMPSUL) and tigecycline (TGC) were utilized for further testing of isolates with concurrent resistance to CIP and AZM. Additionally, an e-test strip for azithromycin was used to further evaluate AZM non susceptibility. This panel of antibiotics tested exceeds the spectrum of the National Antimicrobial Resistance Monitoring System for Enteric Bacteria (NARMS). Zone diameter breakpoints (mm) for *Campylobacter* spp. from the Clinical & Laboratory Standards Institute (CLSI M45) were applied to assess CIP, ERY, AZM and TET resistance. The CLSI zone diameter breakpoints (mm) for *Enterobacteriaceae* were used for GEN, AMC, AMP, CHL and IMP, CLI, FOSF and AMP given that there are no established breakpoints for *Campylobacter* (**Supplementary Table 1**).

## Results and Discussion

A total of 235 *Campylobacter* isolates from 228 fecal samples were concurrently resistant to AZM and CIP. Of these, 111 animal derived isolates and 67 human derived isolates were determined to be *C. coli*. Additionally, 14 human derived isolates and 43 animal derived isolates were determined to be *C. jejuni*. A single human sample and 6 chicken derived samples were determined to have a co-infection of *C. jejuni* and *C. coli* and thus both isolates were included in the analysis. Resistance patterns are shown separately shown for animal and human samples, as well as *C. jejuni* and *C. coli* isolates in **Table 1**. Briefly, 169/178 *C. coli* and 54/57 *C. jejuni* had an e-strip result available. All isolates showed resistance to AZM (n=8 between >16 ug/mL-75 ug/mL; n=215 >256 ug/mL). Resistance to antibiotics included in the extended panel (CLI, FOSF, AMPSUL and TGC) varied. Notably, over 85% of *C. jejuni* and almost 100% of *C. coli* isolates were resistant to CLI, over 65% of both species were resistant to AMPSUL and approximately 30% of both species were resistant to FOSF.

**Table 1.** The distribution of AZM and CIP resistant *Campylobacter jejuni* and *Campylobacter coli* strains derived from human and animal fecal samples.

| Antibiotic | Human Isolates (n=81) |  |  |  |  |  | Animal Isolates (n=154) |  |  |  |  |  |
| --- | --- | --- | --- | --- | --- | --- | --- | --- | --- | --- | --- | --- |
|  | *C. jejuni* (n=14) |  |  | *C. coli* (n=67) |  |  | *C. jejuni* (n=43) |  |  | *C. coli* (n=111) |  |  |
|  | R (%) | I (%) | S (%) | R (%) | I (%) | S (%) | R (%) | I (%) | S (%) | R (%) | I (%) | S (%) |
| Erythromycin (15 µg) | 100.0 | 0.0 | 0.0 | 98.5 | 0.0 | 1.5 | 100.0 | 0.0 | 0.0 | 100.0 | 0.0 | 0.0 |
| Tetracycline (30 µg) | 92.9 | 0.0 | 7.1 | 95.5 | 1.5 | 3.0 | 100.0 | 0.0 | 0.0 | 92.7 | 0.0 | 7.2 |
| Amoxicillin and Clavulanic Acid (30 µg) | 0.0 | 0.0 | 100.0 | 0.0 | 4.5 | 95.5 | 0.0 | 2.3 | 97.7 | 5.4 | 10.8 | 83.8 |
| Gentamicin (10 µg) | 35.7 | 0.0 | 64.3 | 70.1 | 0.0 | 29.9 | 60.5 | 0.0 | 39.5 | 48.6 | 0.9 | 50.5 |
| Chloramphenicol (30 µg) | 0.0 | 0.0 | 100.0 | 7.6 | 1.5 | 90.9 | 4.7 | 2.3 | 93.0 | 7.2 | 4.5 | 88.2 |
| Imipenem (10 µg) | 0.0 | 0.0 | 100.0 | 0.0 | 0.0 | 100.0 | 0.0 | 0.0 | 100.0 | 0.0 | 0.0 | 100.0 |
| Clindamycin (2 µg) | 85.7 | 0.0 | 14.3 | 100.0 | 0.0 | 0.0 | 97.7 | 0.0 | 4.7 | 91.8 | 0.0 | 8.2 |
| Ampicillin Sulbactam (10 µg) | 64.3 | 14.3 | 21.4 | 85.1 | 1.5 | 13.4 | 69.8 | 9.3 | 20.9 | 58.7 | 4.6 | 36.7 |
| Fosfomycin (200 µg) | 28.6 | 7.1 | 64.3 | 44.8 | 3.0 | 52.2 | 30.2 | 4.7 | 65.1 | 34.3 | 0.0 | 65.7 |
| Tigecycline (15 µg) | 0.0 | 0.0 | 100.0 | 1.5 | 0.0 | 98.5 | 2.4 | 0.0 | 97.6 | 0.9 | 0.0 | 99.1 |

The expanded set of antibiotics tested was done to evaluate alternative antimicrobials, particularly oral regimes, for the most common cause of bacterial gastroenteritis. Prior studies of fluroquinolone resistant strains identified fosfomycin, amoxicillin and clavulanic acid and clindamycin as alternatives.^26^ Fosfomycin has been used for the treatment of infectious gastroenteritis for other MDR causes of enterocolitis including *Shigella sonnei*, ^27^ including in clinical trials of *Campylobacter* gastroenteritis.^28^ Cross resistance to other antimicrobials is nominal, and it would theoretically be an option for treating drug resistant dysentery in contexts where rapid and complete phonotypic and genotypic testing may not be rapidly available. Unfortunately, human isolates revealed too high a prevalence of resistance to make this a feasible strategy in this setting. Clindamycin is less appealing as it fails to simultaneously treat other principal causes of bacterial enteritis and was also shown to have a high prevalence of non-susceptibility. Amoxicillin-clavulanate has consistently been identified in *in vitro* testing of resistant clinical isolates of *Campylobacter* as a feasible oral option,^29,30^ although clinical efficacy has not been clearly demonstrated and there has been two cases of clearly described treatment failure ^31^ of patients with isolates that displayed *in vitro* susceptibility. The discrepancy of *in vitro* results between amoxicillin and clavulanic acid and ampicillin sulbactam is notable and likely the result of the β-lactamase enzyme *bla*_OXA-61_ relative resistance to sulbactam as compared with clavulanic acid.^32^ Clinical trials will be required to demonstrate the clinical efficacy of Amoxicillin-clavulanate acid, the only oral antimicrobial agent that had an acceptable antimicrobial susceptibility profile, as there are numerous results in the literature for enteric diseases in which agents that demonstrate in vitro susceptibility are found to have significant rates of failure in the treatment of clinical disease.^33,34^

Fortunately, multi drug resistance *Campylobacter* has typically been sensitive to carbapenems, with non-susceptibility rates of <1% consistent with our finding of no non-susceptibility in this study.^35^ We tested tigecycline as an alternative agent for highly resistant gram negative infections, and also found nearly universal susceptibility (<2%). Although these findings further clarify options for patients with life threatening illnesses, oral options require further evaluation for the management of patients with moderate to severe diarrhea in populations in endemic high burden settings and for large outbreaks that make it challenging to rely solely on injectable agents.

## Figures and Tables

**Supplementary Table 1.** Breakpoints for disk diffusion testing for Campylobacter.

| Antimicrobial Agent | Abbreviation | Resistant (mm) | Intermediate (mm) | Susceptible (mm) | Source |
| --- | --- | --- | --- | --- | --- |
| **Quinolone** |  |  |  |  |  |
| Ciprofloxacin (5 µg) | CIP | ≤20 | 20-24 | ≥24 | a |
| **Macrolide** |  |  |  |  |  |
| Erythromycin / Azithromycin (15 µg) | ERY / AZM | ≤12 | 13-15 | ≥16 | a |
| **Tetracycline** |  |  |  |  |  |
| Tetracycline (30 µg) | TE | ≤22 | 23-25 | ≥26 | a |
| **Aminoglycoside** |  |  |  |  |  |
| Gentamicin (10 µg) | GM | ≤12 | 13-14 | ≥15 | b |
| **Beta lactam** |  |  |  |  |  |
| Amoxicillin + Clavulanic Acid (30 µg) | AMC | ≤13 | 14-17 | ≥18 | b |
| Ampicillin Sulbactam | AMPSUL | ≤11 | 12-14 | ≥15 | b |
| Imipenem (10 µg) | IMI | ≤19 | 20-22 | ≥23 | b |
| **Lincosamide** |  |  |  |  |  |
| Clindamycin (2 µg) | CLI | ≤12 | 13-17 | ≥18 | b |
| **Other** |  |  |  |  |  |
| Chloramphenicol (30 µg) | CHL | ≤12 | 13-17 | ≥18 | b |
| Fosfomycin (200 µg) | FOSF | ≤12 | 13-15 | ≥16 | b |
| Tigecycline (15 µg) | TIGE | ≤14 | 15-18 | ≥19 | b |

a. CLSI. (2016). Methods for Antimicrobial Dilution and Disk Susceptibility Testing of Infrequently Isolated or Fastidious Bacteria. In CLSI guideline M45 (3rd ed.). Clinical and Laboratory Standards Institute.
b. CLSI. (2016). Performance Standards for Antimicrobial Susceptibility Testing. In CLSI supplement M100 (27th ed.). Clinical and Laboratory Standards Institute.

